# From Variability to Synchrony: Non-linear Development of Auditory Neural Responses During the First Year of Life

**DOI:** 10.64898/2026.02.20.706969

**Authors:** Eva Reisenberger, Manuel Schabus, Cristina Florea, Monika Angerer, Michaela Reimann-Ayiköz, Jasmin Preiß, Dietmar Roehm, Dominik P. J. Heib, Claudius Fazelnia, Mohamed S. Ameen

**Affiliations:** Centre for Cognitive Neuroscience Salzburg (CCNS), University of Salzburg, Salzburg, Austria; Research Group Neurobiology of Language, Department of Linguistics, University of Salzburg, Salzburg Austria; Laboratory for Sleep, Cognition and Consciousness Research, Department of Psychology, University of Salzburg, Salzburg, Austria; Department of Geriatric Medicine, Salzburg University Hospital Christian-Doppler-Klinik, Salzburg, Austria; Department of Obstetrics and Gynecology, Salzburg University Hospital Landeskrankenhaus, Salzburg, Austria

**Keywords:** electroencephalography (EEG), infants, brain development, event-related potentials (ERPs), inter-trial phase coherence (ITPC), auditory processing

## Abstract

In humans, the first year of life is characterized by rapid developmental changes, including substantial brain maturation. As a result, neural responses to auditory stimuli undergo marked changes during this period. In this study, we followed 69 infants across their first year of life and recorded high-density electroencephalography (hdEEG) at 2 weeks, 6 months, and 12 months postpartum. Infants were presented with pure beep tones to examine the development of neural responses to auditory stimulation. We analysed event-related potentials (ERPs), inter-trial phase coherence (ITPC), and time-frequency (TF) responses to the beep tones and controlled for arousal state during stimulus presentation. We found that with increasing age, neural responses became more pronounced and showed reduced trial-to-trial variability. Phase synchronization increased from 2 weeks to later developmental stages in a broad low-frequency range (0 to 11 Hz), indicating improved temporal alignment of brain responses over time. However, phase synchronization decreased from 6 to 12 months, suggesting a developmental transition towards more differentiated brain activity. Taken together, these findings demonstrate that auditory maturation during the first year of life follows a non-linear trajectory driven by dynamic changes in neural synchronization, reflecting the progressive refinement of functional neural circuits. Our results thus provide a critical benchmark for understanding the neural dynamics underlying sensory development during this period.

## 1. Introduction

The first year of life involves rapid and extensive brain maturation. During this time, increases in myelination, synaptic density, and neural connectivity lay the groundwork for developing cognitive and perceptual skills (Huttenlocher, 1979; Huttenlocher & Dabholkar, 1997; Moore & Linthicum, 2007; Picton & Taylor, 2007; Tierney & Nelson, 2009). Already during the first months of life, specialized auditory networks enable infants to process speech, detect phonetic changes, or demonstrate sensitivity to familiar voices, for instance (e.g. Dehaene-Lambertz, 2000; Dehaene-Lambertz et al., 2002, 2010; Dehaene-Lambertz & Dehaene, 1994; Gervain et al., 2008; Mehler et al., 1978; Peña et al., 2003; Perani et al., 2011). A well-known developmental trajectory is that, at birth, infants can detect a broad range of speech sounds and, within the first year, their brains begin to specialize in the phonemes of their native language(s), illustrating early neural specialization (Ortiz-Mantilla et al., 2016; Werker & Hensch, 2015; Werker & Tees, 1984). Therefore, understanding how the infant’s brain processes auditory input during this sensitive period is essential for grasping the foundations of sensory and cognitive development.

Prior research has shown that maturational neural changes and emerging organisational patterns influence auditory evoked potentials (AEPs) in infants, affecting both their timing and waveform morphology (Chen et al., 2023; Choudhury & Benasich, 2011; Coch & Gullick, 2011; Edgar et al., 2015; Kurtzberg et al., 1984; Novak et al., 1989; Ohlrich et al., 1978; Picton & Taylor, 2007; Thierry, 2005). Especially early in infancy, AEPs are highly variable (Picton & Taylor, 2007; Taylor & Baldeweg, 2002). Over the course of development, responses evolve, with latencies of AEPs generally decreasing (Kurtzberg et al., 1988; Kushnerenko et al., 2002; Picton & Taylor, 2007; Taylor & Baldeweg, 2002). Developmental changes in amplitude vary and depend on the specific component (e.g. Kurtzberg et al., 1988; Kushnerenko et al., 2002; Kushnerenko et al., 2013; Picton & Taylor, 2007; Taylor & Baldeweg, 2002).

An additional layer of complexity arises when considering the influence of sleep states in this context, as neural and physiological responses can vary across sleep states, including differences in latency and amplitude of AEPs (e.g. Duclaux et al., 1991; Ellingson et al., 1974; Goto et al., 2000; McNamara et al., 2002; Trinder et al., 1990). Although the possible variations in neural responses between sleep states and wakefulness are sometimes acknowledged (e.g. Barnet, 1975; Edgar et al., 2015), a common approach has been to group sleep stages, often referencing early work like Ellingson et al. (1974), who reported similar AEPs during wakefulness and active sleep in a sample of six full-term newborns.

Previous research on the development of sensory processing in infants has largely focused on event-related potentials (ERPs), emphasizing time-locked evoked responses. However, auditory stimulation is known to also drive systematic changes in ongoing neural oscillations and neural synchronization (Klimesch et al., 1998, 2007; Makeig et al., 2002; Sayers et al., 1974). These oscillatory and synchrony-related processes are essential mechanisms supporting cognitive and sensory processing (e.g. Bastiaansen et al., 2002; Buzsáki & Draguhn, 2004; Hendry et al., 2025; Klimesch, 1999; Schack et al., 2002). As ERP analyses are limited in their ability to capture such dynamics, studies that rely solely on evoked responses may overlook associated changes in ongoing neural dynamics, like oscillatory activity, which have also been linked to auditory processing in infancy (e.g. Fujioka et al., 2011; Musacchia et al., 2017; Ortiz-Barajas et al., 2023). Combining ERP and oscillatory measures therefore may offer complementary insights into the maturation of auditory networks.

In this longitudinal study, we aimed to characterize how auditory responses in the brain evolve during the first year of life. Combining ERP and oscillatory analyses, we measured brain responses elicited by pure beep tones in the same group of infants at 2 weeks (2W), 6 months (6M), and 12 months (12M) of age. Data collected 2W after birth, the only measurement point at which sleep data were available, were sleep staged to differentiate between wake and sleep states. This approach allowed us to control for arousal state. We examined ERPs, phase synchronization (as measured by inter-trial phase coherence, ITPC; Tallon-Baudry et al., 1996), and spectral power (as measured by time-frequency analysis, TF) both within and across developmental stages. By doing so, we sought to gain a more comprehensive understanding of the evolving neural dynamics of auditory processing over the course of the first year of life.

## 2. Results

To investigate how infants’ brains respond to auditory stimuli across development, we first confirmed the presence and quality of the auditory-evoked responses on an individual level as well as at each developmental stage. Subsequently, we directly compared neural responses across age groups. All analyses focused on the fronto-central region (see Methods *section 4.6*. for details).

### 2.1. Response Validation on the Individual Level

Preprocessed data showed robust evoked neural responses at the single trial level, confirming the reliability of auditory responses for each subject at each measurement time point. Figures 1A-C illustrate trial-by-trial responses to beep tones from a single representative subject in each age group (A: 2W; B: 6M; C: 12M), demonstrating that the beep tones reliably elicited brain responses. This pattern was clearly observable in all of our participants.

**Figure 1.**
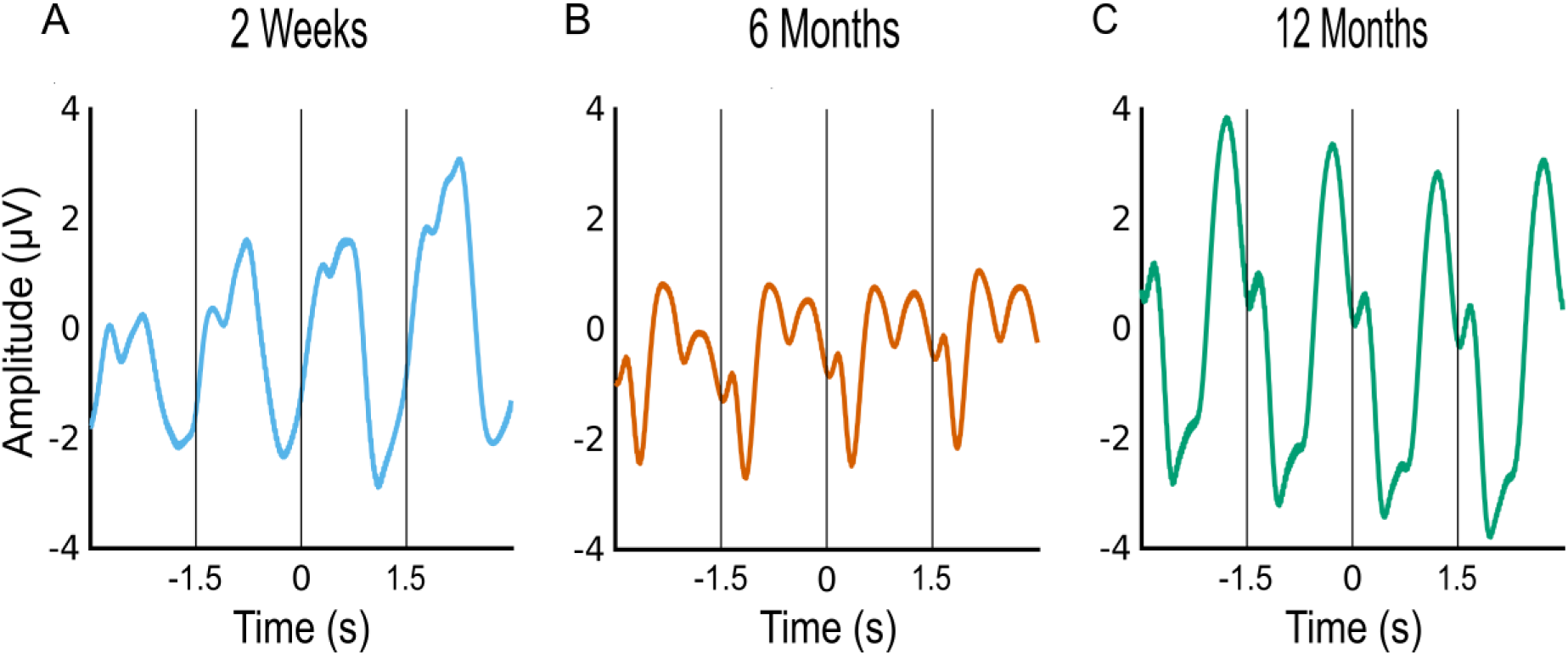
ERP Responses From a Single Subject to Sequentially Presented Beep Tones Measured at A) 2 Weeks (Blue), B) at 6 Months (Orange) and C) at 12 Months (Teal) After Birth. *Note*. Mean event-related potential (ERP) amplitude (µV) is plotted over time (s) in each panel. Each panel (A-C) includes a black vertical line every 1.5 s, indicating the presentation of a beep tone.

### 2.2. Response Validation Across Timepoints

We then characterized the auditory response across all available participants within each age group by comparing brain responses to beep tones against resting brain activity during the no-stimulation condition. The results shown in Figure 2 reflect the averaged neural responses to beep tones alongside the no-stimulation control condition from all available participants in each age group (2W: *n* = 41, 19 female; 6M: *n* = 58, 28 female; 12M: *n* = 53, 25 female). Note that for the 2W measurement point, all available participants, both sleeping and awake, were included in this analysis and that the presented datasets are not directly comparable across ages, as they do not constitute a longitudinal analysis.

**Figure 2.**
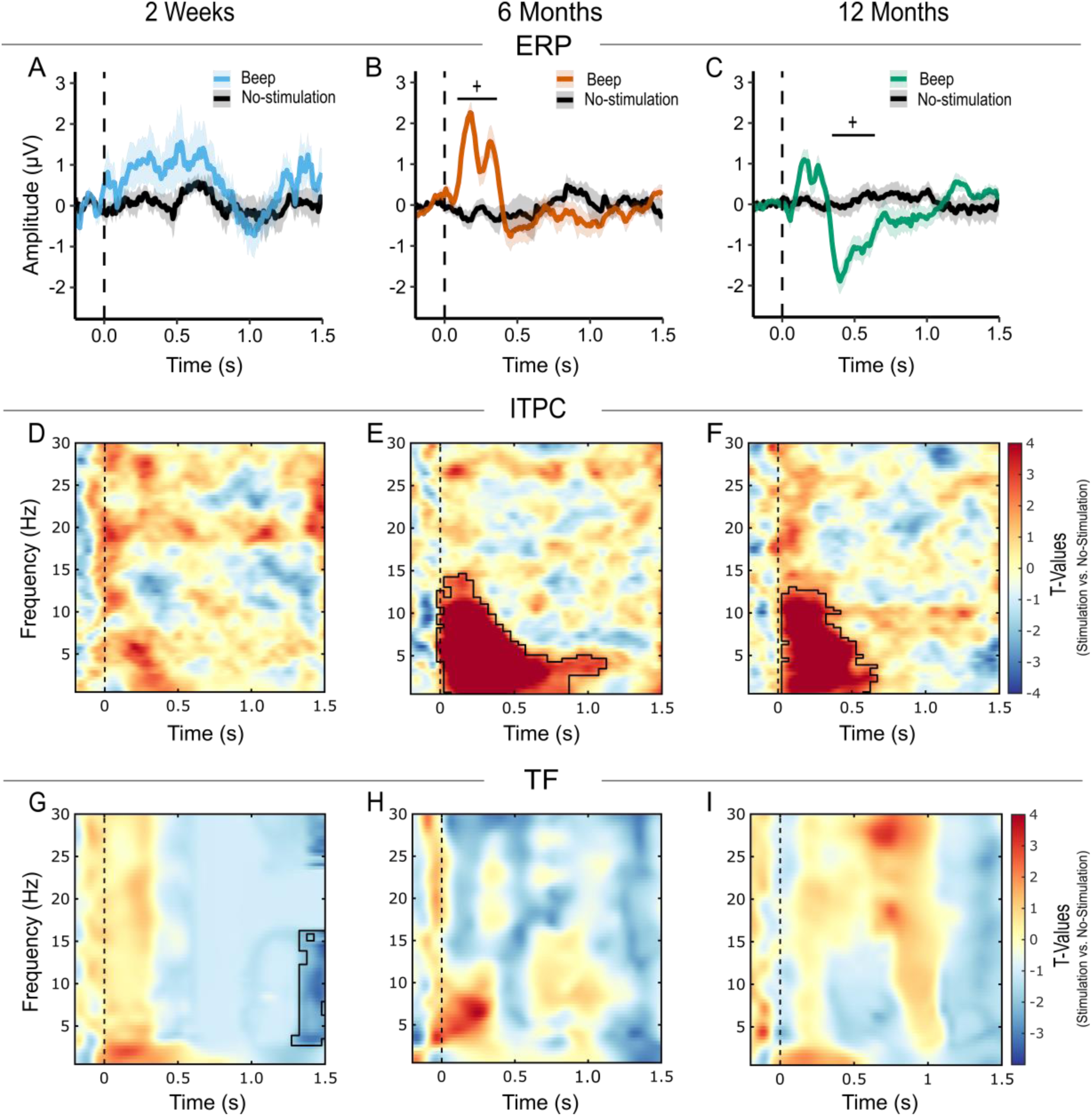
Overview of Auditory Neural Responses Within Age Groups Across Conditions (Stimulation vs. No-Stimulation). *Note.* The signal reflects group level results from all participants available at each measurement time point (2W: *n* = 41; 6M: *n* = 58; 12M: *n* = 53). Columns represent the three measurement time points. Each row displays a different type of analysis: **A-C**) Grand average event-related potential (ERP) waveforms for the stimulation condition at 2 Weeks (blue), 6 Months (orange), and 12 Months (teal). The no-stimulation control condition is plotted in black. Each panel displays mean ERP amplitude (µV) over time (s), aligned to stimulus onset (dashed vertical line at 0s). Shaded regions represent the standard error of the mean. Asterisks indicate statistical significance (\**p* < 0.05, ^+^*p* < 0.01) with the vertical line marking the corresponding time window. **D-F**) Time-frequency (TF) plots depicting *t*-values obtained through cluster-based permutation tests of inter-trial phase coherence (ITPC), representing the difference between stimulation and no-stimulation condition, and **G-I**) TF plots of spectral power differences shown as *t*-values from cluster-based permutation tests comparing the stimulation to the no-stimulation condition. In the second and third row, the TF plots display data across frequencies (0.5 to 30 Hz) and time (-0.2 s to 1.5 s), time-locked to the onset of the stimulus onset (vertical dashed line at 0 s). The color scale represents *t*-values, where positive values (warm colors) indicate higher phase synchronization or spectral power during the presentation of the stimuli, and negative values (cool colors) indicate lower phase synchronization or spectral power relative to the no-stimulation condition. Significant clusters are outlined with a black contour within the TF plots.

For the 2W-old infants (*n* = 41), ERP amplitudes did not differ significantly from the no-stimulation condition (Figure 2A). Among the 6M-old infants (*n* = 58), ERP amplitudes showed significant deviations from the no-stimulation condition, with a significant positive cluster (*p* < 0.001, ∑*t* = 146.33) being observed between 88 ms and 360 ms post-stimulus (Figure 2B), reflecting increased amplitudes during stimulus presentation compared to the no-stimulation condition within the identified time-window. Compared to the no-stimulation condition, the auditory evoked responses of 12M-old infants (*n* = 53) differed significantly as well. The comparison revealed a significant negative cluster (*p* < 0.025, ∑*t* = -122.68) between 344 ms and 640 ms post-stimulus (Figure 2C), reflecting the increased amplitude of the negative ERP component compared to the no-stimulation condition at 12M.

To further explore the source of the ERP amplitude changes, we compared the ITPC between our stimulation and no-stimulation conditions. Among the 2W-olds, no significant clusters in ITPC relative to the no-stimulation condition were observed (Figure 2D). For 6M, ITPC analysis revealed a significant positive cluster (*p* < 0.001, ∑*t* = 1837.79), within an approximately 1100 ms window post-stimulus onset and ranging from approximately 0.5 to 14.50 Hz (Figure 2E). This indicates increased phase synchronization during stimulation relative to the no-stimulation condition across a broad frequency range. Also for 12M a significant positive cluster of increased ITPC was identified (*p* < 0.001, ∑*t* = 1032.67), occurring from approximately 50 ms to 650 ms post-stimulus in the frequency range of approximately 0.50 to 13 Hz (Figure 2F), also reflecting higher phase synchronization in response to stimulation compared to the no-stimulation condition in a broad low-frequency range.

Having established that the auditory beep tones induced changes in both the amplitude and phase of the signal, we next examined whether these sounds also produced alterations in the spectral profile of the neural response. Spectral power analysis revealed a significant negative cluster (*p* < 0.025, ∑*t* = -259.61) for the youngest age group (2W) observed between approximately 1300 ms and 1500 ms post-stimulus, covering frequencies from approximately 3 to 16 Hz (Figure 2G). This indicated reduced power in comparison with the no-stimulation condition within a broad frequency range in the identified time window at 2W. For 6M, spectral power analysis revealed no significant differences in power compared to the no-stimulation condition (Figure 2H). Similarly, for 12M, no significant TF clusters were identified when comparing post-stimulus activity to the no-stimulation condition (Figure 2I).

Overall, auditory responses became more pronounced at later developmental stages, being most clearly observable in the older groups (6M and 12M), where both evoked auditory activity and phase synchronization showed detectable effects. This pattern suggests a developmental strengthening of auditory processing over time. In contrast, spectral power analyses were largely inconclusive.

We also conducted these analyses at temporal sites (see Supplementary Figure A1 in Appendix A). Although the overall pattern of results was comparable across regions, signals recorded at temporal sites were noticeably noisier than those at fronto-central sites. Specifically, in the 2W group, ERP waveforms were less clearly defined, suggesting reduced reliability likely due to a lower signal-to-noise ratio. Therefore, we focused on fronto-central sites in the main analyses.

### 2.3. 2-Week Data Divided By Sleep Stages

To account for potential differences in the infants’ level of arousal during the 2W session, we categorized the data according to four distinct sleep stages: *Wake*, *Active Sleep* (*REM*), *Quiet Sleep* (*NREM*), and *Transitional*, with varying numbers of participants in each group (*n* = 20, 11 female in Wake; *n* = 6, 2 female in REM; *n* = 2, 1 female in NREM; *n* = 6, 3 female in Transitional). Figure 3 depicts the ERP patterns observed in each sleep stage. Given the limited sample sizes, only the Wake group (*n* > 10) could be tested using cluster-based permutation analyses, yielding no significant clusters.

**Figure 3.**
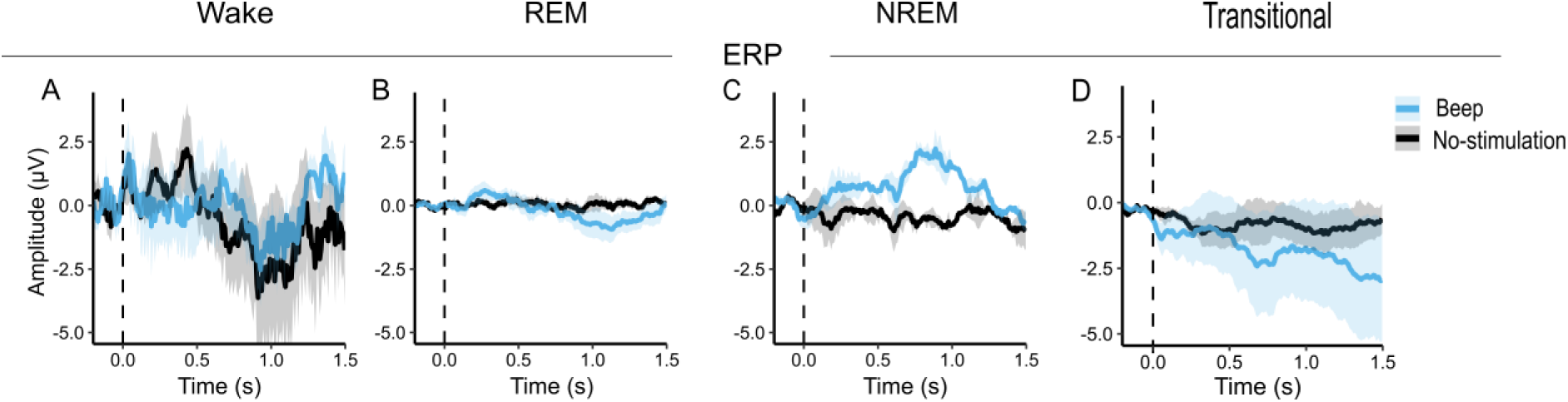
Overview of Grand Average Event-Related Potentials (ERPs) at 2 Weeks After Birth Split by Sleep Stage. *Note*. Columns represent the four different sleep stages (*n* = 20 in Wake; *n* = 6 in REM; *n* = 2 in NREM*; n* = 6 in Transitional). Each panel (A-D) displays grand average ERP waveforms for the stimulation (blue) and no-stimulation (black) condition in these different sleep stages. Shown is the mean ERP amplitude (µV) over time (s), aligned to stimulus onset (dashed vertical line at 0s). Shaded regions represent the standard error of the mean.

Neural responses across these arousal states did not exhibit consistent or robust effects. Corresponding difference plots (stimulation – no-stimulation) of phase synchronization and spectral power are provided in Figure A2 in Appendix A. Note that overall, data across sleep stages and conditions within each sleep stages were available only for small subsets of infants.

### 2.4. Development of Auditory Responses Across the First Year of Life

To examine developmental differences in neural responses to sounds, we compared brain responses to simple beep tones across three age groups (2W, 6M, and 12M), controlling for arousal by including only wake infants. This analysis specifically focused on the same infants measured longitudinally to characterize the developmental trajectory of auditory processing during the first year of life. Sample size varied by pairwise comparison: 2W vs. 6M (*n* = 17, 10 female), 2W vs. 12M (*n* = 15, 8 female), and 6M vs. 12M (*n* = 48, 22 female).

Figures 4A-C display grand-average ERP waveforms in pairwise age-group comparisons. No significant ERP differences were observed for the 2W vs. 6M comparison (Figure 4A), nor for the 2W vs. 12M comparison (Figure 4B). ERP comparison between 6M and 12M revealed a significant negative cluster (*p* < 0.025, ∑t = -69.46) between 296 ms and 432 ms post-stimulus (Figure 4C).

**Figure 4.**
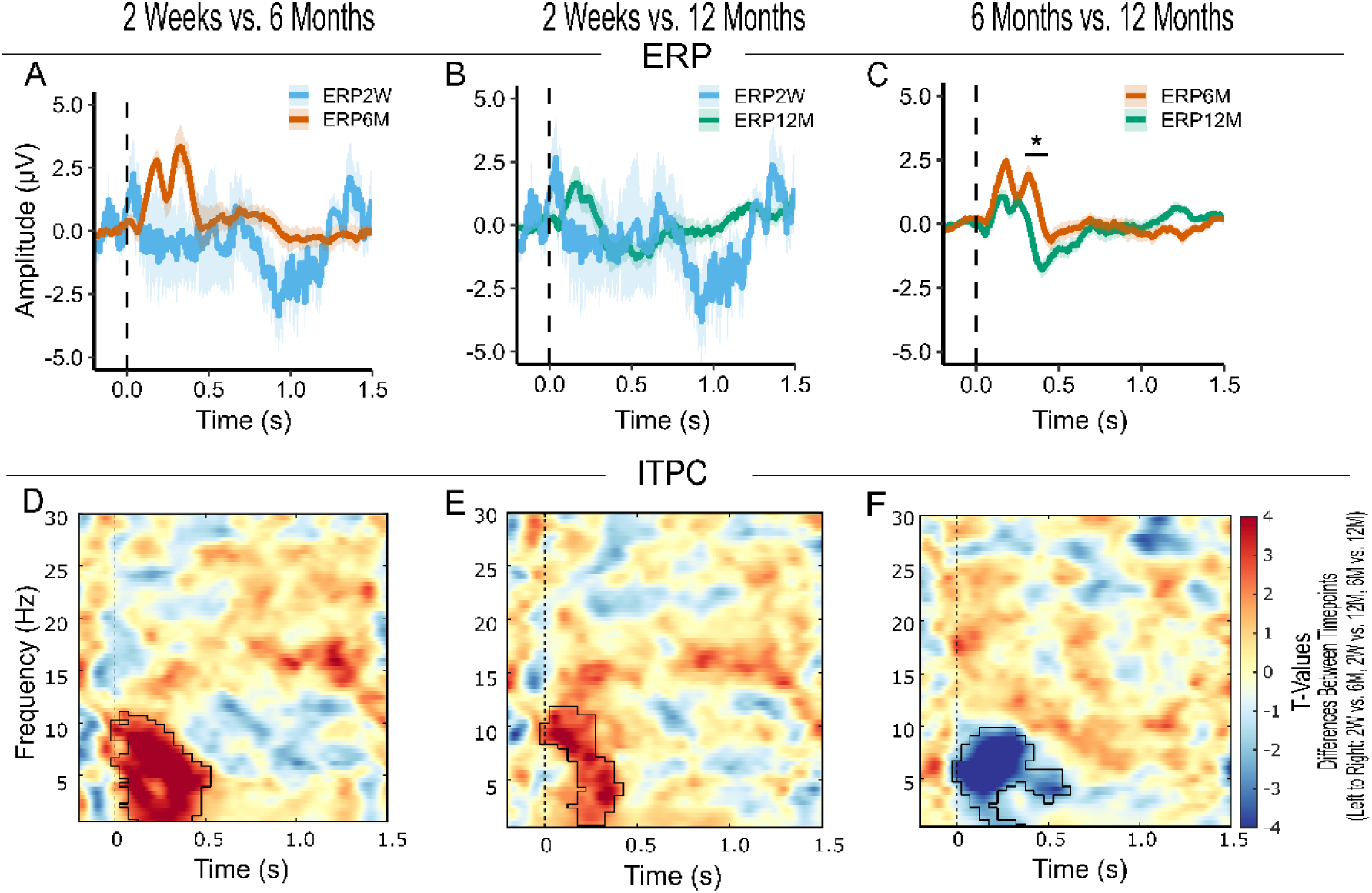
Pairwise Comparisons of Neural Responses to Beep Tones Between Developmental Time Points: 2 Weeks vs. 6 Months; 2 Weeks vs. 12 Months; 6 Months vs. 12 Months. *Note*. Columns represent the three pairwise contrasts: 2 Weeks vs. 6 Months (*n* = 17), 2 Weeks vs. 12 Months (*n* = 15) and 6 Months vs. 12 Months (*n* = 48). Each row displays a different type of analysis: **A-C**) Grand average event-related potential (ERP) waveforms for the stimulation condition at 2 Weeks (blue), 6 Months (orange), and 12 Months (teal). Each panel displays mean ERP amplitude (µV) over time (s), aligned to stimulus onset (dashed vertical line at 0s). Shaded regions represent the standard error of the mean. Asterisks indicate statistical significance (\**p* < 0.05, ^+^*p* < 0.01) with the vertical line marking the corresponding time window. **D-F**) Time-frequency (TF) plots of inter-trial phase coherence (ITPC) showing *t*-values from cluster-based permutation analysis, representing the difference between timepoints (left to right: 2W vs. 6M, 2W vs. 12M, 6M vs. 12M). The TF plots in the second row display data across frequencies (0.5 to 30 Hz) and time (-0.2 to 1.5 s), time-locked to the onset of the stimulus onset (vertical dashed line at 0 s). The color scale for *t*-values represents statistical differences between timepoints, where positive values (warmer colors) indicate higher phase synchronization or spectral power in the older age group, and negative values (cooler colors) indicating higher values in the younger age group respectively. Significant clusters are outlined with a black contour within the TF plots.

Additional statistical analyses of ERP amplitudes and latencies were performed to characterize the developmental trajectory we observed, facilitating comparison with prior studies (e.g. Choudhury & Benasich, 2011; Kurtzberg et al., 1984; Kushnerenko et al., 2002). The negative peak was more pronounced at 12M (*M* = -2.44 µV, *SD* = 2.42 µV) than at 6M (*M* = -1.10 µV, *SD* = 2.66 µV). A paired *t*-test confirmed the change was significant (*t*(47) = 2.91, *p* < 0.01, *d_z_* = 0.42). Moreover, both the first and the second positive peak occurred earlier at 12M than at 6M (first peak at 6M: *M* = 156 ms (*SD* = 47 ms); at 12M: *M* = 134 ms (*SD* = 54 ms); t, *t*(47) = 2.52, *p* < 0.05, *d_z_* = 0.36; second peak at 6M: *M* = 315 ms (*SD* = 41 ms); at 12M: *M* = 274 ms (*SD* = 36 ms); *t*(47) = 4.77, *p* < 0.01, *d_z_* = 0.69).

To further assess developmental changes in neural synchronization, ITPC was compared between the three age groups in pairwise analyses. Figures 4D-F present the corresponding TF representations of *t*-values for ITPC comparisons between age groups. The 2W vs. 6M comparison showed a significant positive cluster (*p* < 0.001, ∑*t* = 604.10), occurring within the approximately 500 ms window following stimulus onset and covering frequencies from approximately 0.50 to 11 Hz (Figure 4D). This indicated an increase in phase synchronization at 6M relative to 2W in a low to mid-frequency range. Similarly, the 2W vs. 12M comparison revealed a significant positive cluster (*p* = 0.001, ∑*t* = 285.92), within an approximately 400 ms window post-stimulus onset and ranging from approximately 1 to 11.50 Hz (Figure 4E). Likewise, this reflects increased phase synchronization across trials at 12M compared to 2W in a broad low-frequency range. In contrast, the 6M vs. 12M comparison revealed a significant negative cluster (*p* < 0.001, ∑*t* = -463.20), within an approximately 600 ms window post-stimulus onset, with a frequency range of approximately 0.50 to 9.50 Hz (Figure 4F). The negative cluster indicates weaker neural phase synchronization across trials at 12M compared to 6M.

Time-frequency analyses using cluster-based permutation tests revealed no significant clusters in pairwise comparisons of age groups. Corresponding figures are provided in the appendix (Supplementary Figure A3 in Appendix A).

Taken together, pairwise comparisons across age groups revealed differences in evoked auditory responses between the two older age groups (6M vs. 12M). Effects in phase synchronization were present for all comparisons, while spectral power analyses showed no significant changes.

## 3. Discussion

Understanding how the infant brain processes auditory stimuli during early development is crucial for mapping the foundations of language and cognition. One of the central challenges in infant EEG research is obtaining sufficiently large and clean datasets while balancing the need for sufficient data with developmental constraints such as attention span limits and immature motor control (Hoehl & Wahl, 2012). By tracking brain responses to simple auditory stimuli in the same infants at 2W, 6M, and 12M of age, we were able to characterize how auditory processing evolves within the first year of life within individuals rather than inferring maturation from cross-sectional comparisons. Our findings highlight a trajectory from immature and highly variable neural responses in the youngest age group to more differentiated ones at later developmental stages, likely reflecting progressive refinement of sensory processing. Overall, the observed patterns suggest that early auditory development reflects progressive refinement and reorganization of neural processing rather than a uniform increase in response magnitude or consistency.

### 3.1. Auditory Evoked Responses and Neural Phase Coherence

First, we examined within-age-group differences between the stimulation and no-stimulation condition. At 2W, the absence of significant differences in auditory evoked responses suggests immature or weak neural responses to the stimuli, as shown by the high variance of the 2W ERPs. By 6M, auditory evoked responses became more robust and better-defined. However, still unstable, the morphology of auditory evoked responses changed further from 6M to 12M as negative amplitudes became more pronounced and dominant, while positive amplitudes were attenuated and their latencies decreased. The observed ERP waveforms at 6M and 12M resemble the previously documented positive-to-negative pattern seen in infancy (Čeponienė et al., 2002; Chen et al., 2023; Choudhury & Benasich, 2011; Kurtzberg et al., 1984; Kushnerenko et al., 2002; Novak et al., 1989; Ohlrich et al., 1978), and the ERPs developed in a way that is consistent with previous research (Chen et al., 2023; Choudhury & Benasich, 2011; Edgar et al., 2015; Kurtzberg et al., 1984; Kushnerenko et al., 2002; Novak et al., 1989; Ohlrich et al., 1978). Complementary ITPC results at 2W showed no significant difference between the stimulation and the no-stimulation condition in phase synchronization, suggesting immature neural phase locking. As compared to 2W-olds, 6M and 12M old infants exhibited enhanced phase synchronization. In comparison to 6M, phase synchronization was observed in a narrower cluster at 12M. These findings suggest a shift towards faster and more efficient processing over the course of development that can be better discussed considering the results of our longitudinal analyses.

For the longitudinal analyses, we leveraged the sleep-scored recordings at 2W and restricted the sample to infants who were awake during stimuli presentation. This ensured that developmental comparisons were not confounded by differences in arousal state across age groups. At the later timepoints (6M and 12M), infants arrived well rested and remained awake throughout the recording. ERP pairwise comparisons between 6M and 12M revealed a significant difference. A latency shift likely is the primary contributor to the observed difference, as earlier peaks were observed in the ERP waveform at 12M compared to 6M. This reduction in peak latencies is in line with prior research showing that faster ERP responses reflect ongoing brain maturation, such as enhanced myelination (Eggermont & Salamy, 1988; Picton & Taylor, 2007). By insulating axons, myelin makes neural signal transmission in infant development faster and more coordinated (Chevalier et al., 2015; Grotheer et al., 2022; Picton & Taylor, 2007). In contrast, we observed no statistically significant differences in auditory evoked responses between 2W and 6M or between 2W and 12M. While this may, at least partly, be due to limited statistical power, this pattern may also reflect genuine developmental characteristics of early auditory processing.

In particular, the maturation of auditory responses during the first year of life may therefore involve refinement of neural timing and synchronization rather than large changes in amplitudes. This interpretation is consistent with our findings of developmental changes in phase synchronization, suggesting that maturation involves qualitative changes in neural dynamics rather than uniform increases in evoked response amplitudes. The observed increases in phase synchronization from 2W to both 6M and 12M in the approximately 1 to 11 Hz range support the observation that phase-locking mechanisms in general become increasingly temporally consistent with age. This likely also reflects ongoing brain maturation. Both myelination and synaptogenesis influence how stimuli are processed. Myelination supports and accelerates the transmission of neural signals (Chevalier et al., 2015; Grotheer et al., 2022; Picton & Taylor, 2007). At the same time, synaptogenesis rapidly increases the number of synapses during the first year in parallel with dendritic and axonal growth (Huttenlocher, 1979; Huttenlocher & Dabholkar, 1997). Together, these processes support a transition from broadly synchronized neural activity in early infancy to enhanced communication across brain regions and increasing circuit specialization (Bosch-Bayard et al., 2022; Falivene et al., 2024; Fujioka et al., 2011; Perani et al., 2011).

Interestingly, however, the comparison between 6M and 12M revealed a significant decrease in phase synchronization at 12M compared to 6M (in the range from approximately 0.5 Hz to 9.50 Hz). This could be related to a more function-specific tuning of neural responses during the first year of life. Newborns and young infants in the first half-year of life show sensitivity to rhythmic patterns in musical and linguistic input (Hannon & Trehub, 2005a, 2005b; Mehler et al., 1988; Nazzi et al., 2000; Telkemeyer et al., 2009; Winkler et al., 2009) and infants’ neural activity responds selectively to rhythmic auditory input (Cirelli et al., 2016; Fujioka et al., 2011). By the age of 12M, however, this broad sensitivity towards rhythmical structures declines (Hannon & Trehub, 2005b), and cortical rhythmic activity becomes more specialized (Fujioka et al., 2011). This reflects the tuning of auditory and cognitive systems to relevant environmental input, consistent with the perceptual narrowing for speech and music (Hannon & Trehub, 2005b; Kuhl et al., 2006; Werker & Tees, 1984). From this perspective, when a stimulus is perceived as salient (because it is rhythmic) and elicits reliable time-locked neural responses, repeated presentations could result in high phase coherence as is the case in our sample at 6M of age. Developmentally, the observed following decrease in phase coherence between 6M and 12M does not mean that infants’ brains are getting worse in processing auditory input. Instead, it could mean that at 12M the brain relies less on externally imposed timing, which reduces strict phase locking to repetitive stimuli.

This idea could also be linked to another concept that is a bit more far-fetched and cannot yet be directly linked to infant development but could nevertheless be brought into play here: sensory predictions. Repeated sensory stimuli like the beep tones in our study provide temporal structure for predicting event timing, and anticipatory sensory events can reset the phase of low-frequency oscillations, allowing neurons to reach peak excitability at the expected moment (Arnal & Giraud, 2012). The observed decrease in phase coherence from 6M to 12M in a lower frequency range (up to 10 Hz) could also reflect a corresponding developmental shift from reliance on temporal regularities toward more flexible, content-driven predictive strategies. Although this interpretation is highly speculative, it suggests a potentially interesting direction for studying how predictions mature in infancy.

Considering the developmental dynamics of phase coherence in our data, a matching pattern has previously been observed for speech stimuli, where 6M old infants showed stronger 2-6 Hz phase coherence than 12M-olds, alongside higher high-frequency (> 70 Hz) power in the younger age group (Ortiz-Mantilla et al., 2016). These findings have been interpreted as reflecting increasingly refined and efficient neural processing (Ortiz-Mantilla et al., 2016). Considering all of the above, we speculate that development may be less about uniformly increasing phase coherence but more about using coherence selectively and that our observed decrease between 6M and 12M therefore also reflects increasing efficiency.

Accordingly, our ITPC results further support the view that early auditory development reflects progressive refinement of neural processing dynamics, rather than a simple linear increase in brain responses to sounds. Importantly, we think that the two perspectives we brought up (i.e., brain maturation and specialization) go hand in hand. The increase in ITPC from 2W to 6M and 12M could be explained by structural improvements in neural timing and connectivity, while the decrease between 6M and 12M may reflect refinement and the emergence of predictive, and linguistically relevant processing. Together they help to develop an understanding of non-linear developmental trajectories of phase coherence across the first year of life.

We also want to note here that while we interpret our findings as reflecting developmental changes in auditory processing, this interpretation should be considered with caution. The observed effects may not be exclusively auditory, as factors such as attention and sensitivity to rhythmic structure can influence neural responses to auditory stimuli. These processes undergo substantial maturation during the first year of life and are closely linked to the development of auditory cortical function (e.g Cirelli et al., 2016; Fujioka et al., 2011; Hannon & Trehub, 2005a, 2005b; Mehler et al., 1988; Nazzi et al., 2000; Telkemeyer et al., 2009; Winkler et al., 2009). Therefore, our findings likely reflect not only the maturation of auditory pathways but also the progressive refinement of neural systems supporting temporal coordination and sensory processing more broadly.

We further note that all ITPC effects we observed fell within a lower frequency range (up to 13 Hz), which has been shown to be robustly modulated by diverse auditory stimuli (e.g. speech, tones, clicks) in adults (e.g. Hsiao et al., 2009; Krause et al., 1997; Luo & Poeppel, 2007) and infants (e.g. Attaheri et al., 2022, 2024; Carr et al., 2021; Kalashnikova et al., 2018; Kaminska et al., 2025; Musacchia et al., 2015; Ortiz-Barajas et al., 2023; Ortiz-Mantilla et al., 2016). At this point, it is also worth noting that we deliberately chose not to categorize the observed frequency ranges into conventional band labels such as *Theta* or *Alpha*. While these labels are commonly used in adult EEG research, their definitions are not well established in the context of infancy and vary across developmental EEG studies. Some are reusing adult labels (e.g. Attaheri et al., 2024 – Theta: 4 to 8 Hz, Alpha: 8 to 12 Hz) while others introduce infant-specific ranges (e.g. Xie et al., 2018 – Theta: 2 to 6 Hz, Alpha: 6 to 9 Hz). These inconsistent criteria for defining frequency bands in infant studies may introduce interpretive challenges. To maintain clarity and facilitate reproducibility, we reported the actual frequency ranges associated with significant effects.

### 3.2. Spectral Power Results

Spectral power analyses within and across age groups were largely inconclusive. The only observable effect was a late decrease in power at 2W for the stimulation condition compared to the no-stimulation condition (between 1300 ms and 1500 ms after stimulus onset, ranging from approximately 3 to 16 Hz). At first glance, this might suggest a desynchronization pattern prior to the next stimulus. However, no decrease was visible in the pre-stimulus window (-200 ms to 0 ms), making it unlikely to reflect anticipatory desynchronization. Overall, the interpretation of this effect in spectral power is not straightforward, and its origin remains unclear.

The observed discrepancy between the ERP analysis and oscillatory measures offers considerable potential for future research looking at different components contributing to the overall electrophysiological signal. In general, the presence of significant effects in auditory evoked responses and phase consistency at 6M and 12M could support the view that auditory stimuli induce phase resetting of ongoing neural oscillations (e.g. Makeig et al., 2002; Sayers et al., 1974). Future studies should investigate the spatial and mechanistic underpinnings of evoked responses to clarify their functional significance. Neural activity has been shown to compromise a notable aperiodic (non-oscillatory) component shaping the electrophysiological signal alongside oscillatory activity (Donoghue et al., 2020; Voytek et al., 2015). Consequently, ERPs may capture both phase-locked oscillatory activity and aperiodic influences (Ameen et al., 2025). This dual contribution may explain why ERP and oscillatory analyses yielded different results but based on our data and analysis, firm conclusions about underlying mechanisms cannot be drawn.

### 3.3. Sleep Stage-Related Variability in Neural Responses at 2W

To account for infants’ varying levels of arousal at 2W, we analyzed neural responses across four sleep stages: *Wake*, *Active Sleep (REM)*, *Quiet Sleep (NREM)*, and *Transitional*. Our exploratory, though underpowered, analysis indicates that arousal state may indeed affect amplitudes and latencies (e.g. Duclaux et al., 1991; Ellingson et al., 1974; Goto et al., 2000; McNamara et al., 2002; Trinder et al., 1990). Future studies should investigate the development of early auditory responses within each sleep stage in a larger dataset.

### 3.4. Choice of Electrodes

Although temporal regions are more associated with auditory responses, ERP analyses at temporal sites did not reveal a reliable signal-to-noise ratio. We attribute this mainly to challenges inherent to infant EEG, such as posture management and head support, particularly in our 2W-old sample (Hoehl & Wahl, 2012). Given that temporal sites are lateral scalp locations, we think they were likely more affected than the fronto-central area.

That is why we selected the fronto-central region as our region of interest. Indeed, most infant ERP studies include fronto-central electrodes as part of the electrode set analysed (Čeponienė et al., 2002; Kushnerenko et al., 2002, 2007; Leppänen et al., 2004; Novak et al., 1989; Novitski et al., 2007; Telkemeyer et al., 2009), some even exclusively focused on the fronto-central region (e.g. Cheour et al., 2002; Katus et al., 2020; Seery et al., 2014; Urbanec et al., 2024). Moreover, prior infant ERP research indicates that several components in newborns are typically maximal over fronto-central sites (Kurtzberg et al., 1984; Novak et al., 1989; Picton & Taylor, 2007; Telkemeyer et al., 2009), supporting a focus on these regions in infant ERP studies. Accordingly, we consider fronto-central sites the most appropriate and sensitive region of interest for the present study. For context, it is worth noting that much of the cited literature here examined various effects and developmental processes (e.g. mismatch responses, maturation of auditory ERPs, and links between ERP markers and neurodevelopment).

### 3.5. Limitations

Our study faces constraints due to a high number of artifacts in the signal, a challenge that is typical for infant research. Already upon visual inspection, the 2W data appeared highly variable and considerably noisier than the 6M and 12M data. As a result, we must always consider that some effects or their absence might be influenced by variability in signal quality.

In terms of analysis, controlling for sleep stage clearly reduced our sample size. Splitting the sample by arousal state was meant to improve interpretability and comparison across different developmental stages but at the same time underscored the challenges and complexity of working with (sleep) EEG data from such young children. For such young infants, sleep staging per se remains challenging even for trained raters. The inherent complexity, combined with the absence of dedicated EOG channels due to time constraints in the 2W EEG-montage, introduced a degree of uncertainty and likely reduced the precision of stage classification as EOG channels are important for detecting eye movements and, for instance, identifying REM sleep. Despite best efforts, manual scoring of sleep stages cannot be considered perfectly and entirely reliable. Consequently, the development of automated sleep staging techniques is warranted, though, to our knowledge, their implementation with infant data is not yet sufficiently developed. So, even though wake-only data improves interpretability and consistency, it comes at the cost of statistical power.

Another limitation relates to the testing environment. Most of the testings conducted 2W after birth (*n* = 34 out of the 41 datasets included in the final analysis) were carried out in the families’ homes. This approach may have introduced additional variability, as the home setting is less controlled compared to a laboratory environment. Factors such as background noise and positioning could affect the quality of the recorded data and potentially influence the reliability of the measures used. While testing in naturalistic settings is generally desirable, its methodological effects remain somewhat unclear and warrant for further investigation.

Moreover, our sample was relatively homogenous, consisting primarily of infants from similar demographic backgrounds. While this likely reduced variability related to environmental and genetic factors, it also constraints generalizability of the results. Early neurodevelopment and auditory processing are shaped by a complex interplay of biological and experiential influences, e.g. nutritional factors (Cheatham, 2020), maternal sleep quality during pregnancy (Lavonius et al., 2020) or perinatal maternal anxiety (Harvison et al., 2009), which we did not control for. Future research with more diverse populations would be desirable and could also provide valuable insights into how specific environmental factors shape early auditory perception.

### 3.6. Conclusion

The aim of this study was to characterize the development of neural responses to auditory stimuli throughout the first year of life. At 2W, auditory responses were highly variable and weak, likely reflecting immaturity. By 6M, more pronounced ERP components and significantly increased ITPC emerged, indicating stronger temporal alignment of neural responses. By 12M, ERP morphology changed further and phase synchronization, even though still higher than at 2W, decreased. Our findings suggest progressive refinement of auditory processing, allowing infants to process auditory stimuli faster and more efficiently by 12M of age. Overall, the directions of the effects observed point toward a non-linear developmental trajectory and generally reflect complex changes in neural responses during the first year of life.

## 4. Methods

### 4.1. Participants

The project followed a cohort of pregnant women and their children from the 34^th^ week of gestation through the first 12 months postpartum. Ethical approval for the study was obtained from the Ethics Committee of the University of Salzburg (EK-GZ: 12/2013).

A total of 72 pregnant women were initially recruited for the study. Three participants withdrew prior to childbirth, resulting in a sample size of 69 mother-infant dyads (32 female infants) at the first recoding point 2W after birth. After this initial recording, two additional participants discontinued participation. Hence, in total 67 mother-infant dyads (31 female infants) successfully completed the full study protocol up to 12M postpartum. Participants were recruited through multiple channels, including local media and collaborations with institutions providing services to pregnant women (e.g. prenatal yoga classes, healthcare providers etc.). All participants enrolled on a voluntary basis. Prior to participation, all mothers received detailed information regarding the study’s aims and procedures and provided written informed consent. Participants were compensated monetarily throughout the study, with amounts determined for each testing session, in addition to receiving small gifts as a gesture of appreciation.

Inclusion criteria required pregnant women to be at least 18 years old, with no upper maternal age limit. They also needed to report German as their native language and intend to raise their child in a predominantly German-speaking environment. Pregnancies classified as high-risk – such as those with anticipated preterm birth at the time of application – or twin pregnancies were excluded. Participant demographics and child-specific information were collected using a structured, study-specific questionnaire. Key sample characteristics of the recruited sample, including maternal demographics (Table B1) as well as birth and newborn characteristics (Table B2) are provided in Appendix B.

The present analysis is based on EEG data collected from the same infant cohort at three distinct developmental stages – 2W, 6M, and 12M postpartum – as illustrated in Figure 5A.

**Figure 5.**
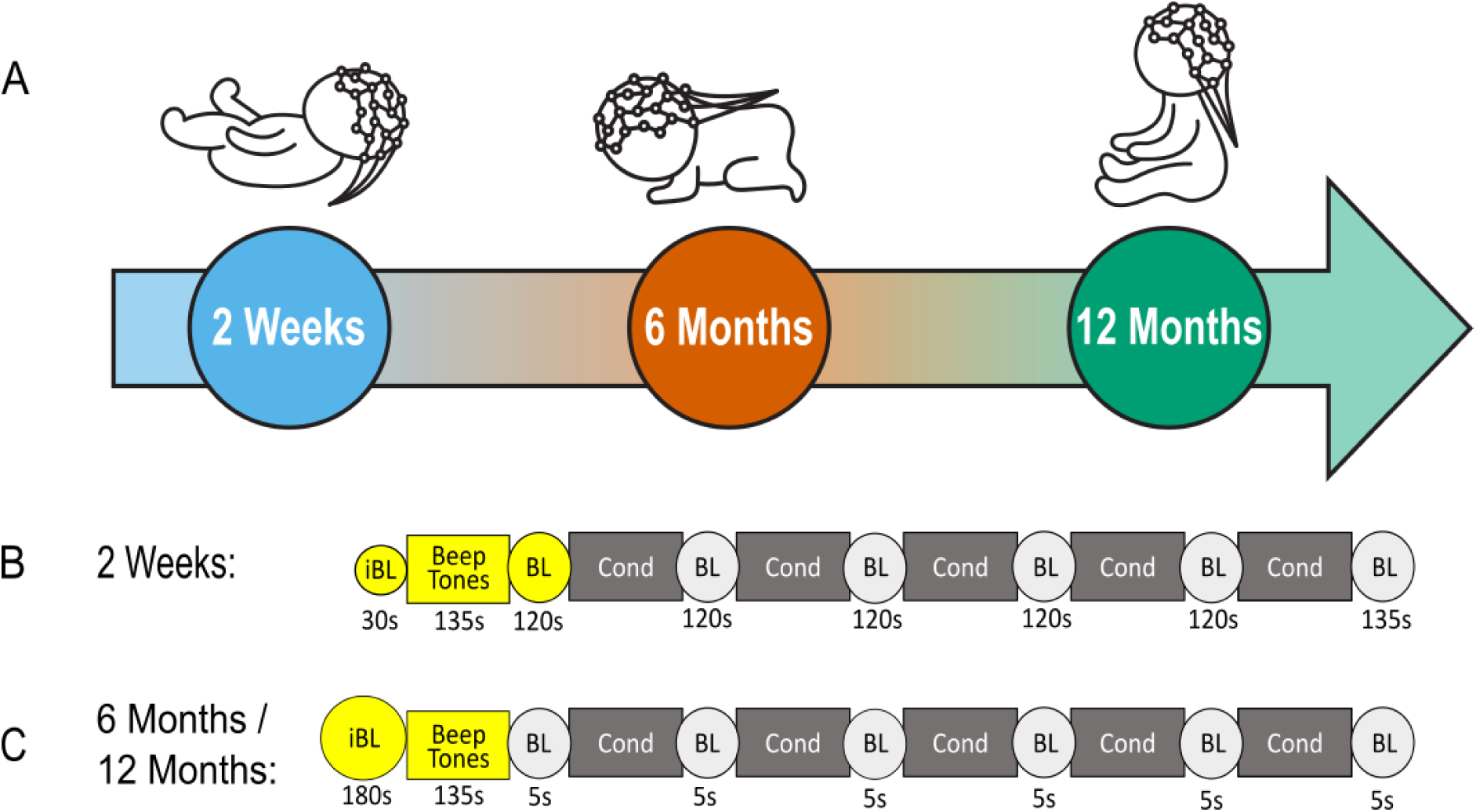
Depiction of the Experimental Design. *Note.* **A**) The different EEG testing timepoints are illustrated with baby icons to aid visualization. Babies were tested at the age of 2 weeks, 6 months, and 12 months in varying positions across developmental stages and participants. Therefore, the infants’ positions depicted in the icons serve illustrative purposes only. See Table B3 in Appendix B for the actual variations in positioning of the infants during the relevant segment of the recording. **B-C**) **Paradigm.** The first testing session 2 Weeks after birth differed from the later ones at 6 Months and 12 Months in the length of the silent initial baseline period (iBL) and the baseline periods without any auditory stimulation (BL) separating experimental conditions (Cond). In the later sessions, the durations of silent baseline periods were decreased to accommodate age-related changes in infants’ attention span and receptiveness. In the present analysis, only the yellow-marked segments were considered for analysis. **B**) **2 Weeks Session**: The testing began with a 30 s silent initial baseline period (iBL), followed by a 135 s auditory stimulation period, during which auditory beep tones were presented (Beep Tones). This was followed by a sequence of alternating baseline periods without auditory stimulation (BL) and experimental conditions (Cond). **C**) **6 Months/ 12 Months Session:** The session started with a 180 s initial baseline period (iBL), followed by a 135 s “Beep Tones” period, during which auditory beep tones were presented (Beep Tones). Thereafter, 5 s silent baseline periods (BL) and experimental conditions (Cond) alternated.

At the first measurement point, conducted approximately 2W after birth, EEG data were collected from 69 infants (*n* = 69, 32 female; *M* = 16.42 days, *SD* = 3.47, Range = 12 to 33). Of these, 28 datasets were excluded from further analyses due to the following reasons: (i) excessive movement and fussiness leading to poor data quality (*n* = 21, for details on the assessment of data quality, see Methods Section *4.6.1*.), (ii) a modification in the experimental paradigm after data collection had commenced, resulting in the removal of four datasets (*n* = 4), (iii) technical issues during the recording (*n* = 1), and (iv) prenatal exposure to maternal smoking (*n* = 1), as previous research has linked smoking during pregnancy to adverse effects on neurodevelopment and altered neurophysiological responses in infants (Kiechl-Kohlendorfer et al., 2010; Shea & Steiner, 2008; Stroud et al., 2018), and (v) another one because the child was already 33 days old on the day of the recording (*n* = 1). This resulted in a final sample of 41 valid datasets for the first time point 2W postpartum (*n* = 41, 19 female; *M* = 15.76 days, *SD* = 2.42, Range = 12 to 25).

At the second measurement point, approximately 6M after birth, EEG data were collected from 67 infants (*n* = 67, 31 female; *M* = 6.16 months, *SD* = 0.21, Range = 5.57 to 6.87). Nine datasets were excluded: (i) Eight due to poor data quality resulting from movement artifacts (*n* = 8), and (ii) one due to continued exclusion of the participant exposed to maternal smoking (*n* = 1). This yielded 58 datasets for analysis at 6M of age (*n* = 58, 28 female; *M* = 6.14 months, *SD* = 0.19, Range = 5.57 to 6.7).

At the 12M follow-up, EEG sessions were conducted again in the same sample of 67 infants (*n* = 67, 31 female; *M* = 12.16 months, *SD* = 0.36, Range = 11.3 to 13.6). Fourteen datasets were excluded for the following reasons: (i) poor data quality due to movement artifacts (*n* = 9), (ii) technical problems during data acquisition (*n* = 3), (iii) continued exclusion of the smoking-exposed participant (*n* = 1) and (iv) inability to record EEG data because the infant refused to wear the cap (*n* = 1). This resulted in a final sample of 53 usable datasets for the last testing session at the age of 12M (*n* = 53, 25 female; *M* = 12.20 months, *SD* = 0.37, Range = 11.3 to 13.6).

### 4.2. Experimental Paradigm

This study is part of a longitudinal research project, which aims to investigate the early foundations of neurodevelopment and to explore how prenatal and postnatal factors shape (cognitive) development and the formation of attachment.

The first part, conducted at 2W after birth, was completed at the families’ homes for 34 out of the 41 dyads included in the final analysis, while the remaining participants (*n* = 7) took part in the same procedure in the laboratory. Most of the follow-up testings used for analysis at 6M (*n* = 58 out of 58) and 12M (*n* = 52 out of 53) took place in the laboratory. Home visits at these later stages were conducted only in exceptional cases (*n* = 1 out of 53 for the third timepoint at 12M). Video was recorded during all EEG sessions.

Upon arrival at the testing site, we measured the infants’ head circumference and fitted them with an EEG cap of appropriate size. At each measurement point, the recording started with a silent baseline period followed by an auditory stimulation sequence consisting of a series of 1000 Hz pure beep tones. This stimulation phase lasted 135 s, during which 90 beep tones were presented at 1.5 s interstimulus intervals, with each tone lasting 100 ms. Following the beep tone sequence, alternating blocks of nursery rhymes and baseline periods without any auditory stimulation (see Figure 5B) were presented. However, only the beep tone sequence, its adjacent silent baseline periods are relevant for the present analysis (see Florea et al. (2024) for further details on the full protocol and analysis of the rhyme paradigm).

#### 4.2.1. Stimulation Procedures

The duration of the silent baseline periods surrounding the tone sequence varied across the three testing time points. At 2W, the session began with a 30 s silent baseline, followed by a 135 s beep tone sequence and a subsequent 120 s baseline period without any auditory stimulation (see Figure 5B). At 6M and 12M, the initial silent baseline period was extended to 180 s, while the silent baseline period following the presentation of the beep tones was reduced to 5 s (see Figure 5C). These adjustments were made to reduce the overall session duration and to accommodate the developmental constraints of attention span and tolerance for experimental procedures in infants. At 2W, we tolerated infants sleeping during the session, as sleep at this age is largely unavoidable, irrespective of the stimulation provided. This allowed for a stimulation duration of approximately 29.5 min. At 6M and 12M infants were awake and alert, and session length was approximately 20 min.

### 4.3. Stimulation protocol

Auditory stimuli were presented at approximately 60 dB sound pressure level (SPL). Before each testing session, we verified the intensity using a calibrated sound pressure level meter with an artificial ear to ensure it remained close to 60 dB, allowing for slight variation. Loudspeakers were positioned one meter away from the infant’s head, though minor deviations in distance could occur as a result of the infant’s movement throughout the session.

To ensure comfort for both the infant and the parent, caregivers were free to choose the most convenient position during data collection. Therefore, the infant’s position was not standardized and varied both across and within sessions, depending on what was most comfortable and feasible in the moment. Infants’ positions also changed with age, from mostly lying at 2W to sitting at 6M and 12M. Detailed information on the positioning of the infants during the beep tones sequences specifically is provided in Table B3 in the Appendix. Caregivers were instructed to keep their child as still as possible. If the infant became fuzzy, the caregiver and/or experimenter used various calming strategies (breastfeeding, offering food or a pacifier, using toys, or blowing soap bubbles) to keep the infant calm and comfortable. These calming strategies were kept to a minimum to avoid overstimulation.

To prevent maternal reactions to the auditory stimuli from influencing the infant, mothers wore noise-canceling headphones and listened to a recording of rainfall sounds. At the 2W testing session, two mothers wore standard (non-noise-canceling) headphones instead, and one mother did not hear the rainfall sounds during the beep segment due to technical issues. However, inspection of the according single-subject data revealed no noticeable differences, so their data was retained.

A trained researcher was present in the room during all testing sessions. At the 2W timepoint, additional individuals – such as the infant’s father or support staff – were occasionally present. All individuals in the room were instructed to remain quiet and minimize movement. The infants’ state of arousal varied only at the first timepoint. For the 6M and 12M sessions, families were asked to ensure that the infant was well-rested when arriving to the appointment so that they stayed awake during the whole session.

### 4.4. Data Acquisition

EEG was recorded using a 124/128-channel HydroCel Geodesic Sensor Net (Electrical Geodesic, Inc.// EGI, Inc., Eugene, OR, USA) and a Net Amps 400 amplifier (Electrical Geodesic, Inc.; EB NEURO S.p.A., Firenze, Italy). Before application, the EEG cap was soaked in a warm water solution containing potassium chloride and baby shampoo. Electrode impedances were kept below 50 kΩ whenever possible, and the sampling rate was set at 1000 Hz, using the midline central electrode (Cz) as an online reference.

### 4.5. Sleep Staging

For sleep staging, the EEG files were rated in 30 s epochs. Sleep stage analysis followed the guidelines for children by the American Academy of Sleep Medicine (AASM) sticking to the *AASM Manual Version 2.1* (Berry et al., 2014) and the *AASM Manual 2.2* (Berry et al., 2015). Sleep stages were divided into *Wake*, *Active sleep (equivalent to rapid eye movement [REM] in adults), Quiet sleep (equivalent to non-rapid eye movement sleep [NREM] in adults)*, and *Transitional* stages (Berry et al., 2014, 2015). Classification was based on EEG as well as respiration and movement patterns observed in the recorded video. Sleep scoring was done manually by an independent and trained expert in the field.

### 4.6. EEG Preprocessing and Analysis

EEG data was preprocessed using the EEGLAB toolbox (Delorme & Makeig, 2004; Version 2024.1) and a custom MATLAB (R2023b Update 10 23.2.0.2859533; The MathWorks, Inc., Natick, MA, USA) code. First, physiological channels as well as the two outermost rows of EEG electrodes were excluded due to higher artifact risk and instability, especially when testing 2W-olds who required head support, which could distort electrode contact and placement (Hoehl & Wahl, 2012). Thus, the following preprocessing steps were performed on a total of 91 electrodes. Data were band-pass filtered between 0.1-45 Hz using a finite impulse response (FIR) filter. Next, the PREP-Pipeline was used for bad channel rejection and interpolation and the application of a robust average reference (Bigdely-Shamlo et al., 2015). As the PREP-Pipeline was originally designed for adult EEG, we adjusted the standard thresholds to infant EEG and focused on extreme outliers. A channel was rejected (i) when the power above 50 Hz, relative to power below 50 Hz, had a Z-score greater than 8, (ii) when its overall robust deviation had a Z-score greater than 13, or (iii) if it lacked correlation with other channels in more than 10% of time windows. Artifact removal was performed using independent component analysis (ICA) with automated component classification and rejection via ICLabel (Pion-Tonachini et al., 2019) following the established thresholds for identifying non-brain components (muscle, ocular, heart, line noise and channel noise; all ≥ 0.80 classification probability). For the datasets included in the final analysis (2W: *n* = 41; 6M: *n* = 58; 12M: *n* = 53) component rejection during preprocessing was as follows: 2W: *M* = 4.80, *SD* = 4.53, ranging from 0 to 18; 6M: *M* = 2.62, *SD* = 2.89, ranging from 0 to 15; 12M: *M* = 2.09, *SD* = 2.68, ranging from 0 to 15.

All preprocessing steps that were implemented served to optimize data quality while maintaining compatibility with common and reproducible EEG analysis practices. By implementing each step individually, we were able to systematically inspect the data after each stage – such as filtering, bad channel detection, and artifact removal – to verify the impact of each procedure on data quality.

All analyses were performed in FieldTrip (Oostenveld et al., 2011) and focused on the fronto-central region. Six electrodes often included in infant ERP research (Burden et al., 2007; Isler et al., 2012; Urbanec et al., 2024) were selected based on their relevance to the region of interest (E24 (F3), E124 (F4), E11 (Fz), E36 (C3), E104 (C4), E129 (Cz); see Figure A4 in Appendix A).

#### 4.6.1. Event-related Analysis

A high-pass filter with a cutoff frequency of 0.5 Hz was applied to the EEG data. EEG data were then resampled to 125 Hz and epoched from -200 ms to 1500 ms relative to stimulus (beep tone) onset.

We created a control (no-stimulation) condition by selecting epochs from baseline periods without auditory stimulation, either preceding or following the presentation of beep tones. The no-stimulation epochs were matched to the stimulation epochs in duration (1700 ms) and number of trials to ensure comparability. As in the stimulation condition, no-stimulation epochs were baseline-corrected using the first 200 ms, thereby generating a relative change signal and ensuring consistency in preprocessing across conditions. For the 6M and 12M recordings, the no-stimulation condition was derived from the initial baseline period preceding the presentation of beep tones. Specifically, a 160 s segment was extracted from the 180 s baseline, beginning 10 s after the baseline onset marker and ending at 170 s. This excluded the first and the last 10 s to minimize artifacts related either to the start of the recording or the onset of the stimulation protocol. Epochs were sampled at 1000 ms intervals, resulting in a 700 ms overlap between consecutive epochs. For the 2W recordings, no-stimulation epochs were extracted either from the initial baseline period preceding beep tones presentation (30 s) or from the baseline period following the presentation of beep tones (120 s), depending on the analysis. For longitudinal and sleep-stage-controlled analyses, epochs were extracted from the initial 30 s baseline period (specifically between 8 s and 28 s), ensuring that infants remained in the same arousal state and allowing for consistency across age groups. Epochs were sampled every 200 ms, resulting in a 1500 ms overlap between consecutive epochs. This high degree of overlap was necessary to obtain a sufficient number of epochs for reliable estimation of neural response metrics. For analyses not constrained by arousal state, no-stimulation epochs were extracted from the longer baseline period following the presentation of beep tones to maximize data availability while still avoiding edge artifacts.

Noisy epochs rejection was applied to both no-stimulation and stimulus epochs, removing trials with large amplitude fluctuations exceeding 1000 μV. The number of rejected trials per participant (2W: *n* = 41; 6M: *n* = 58; 12M: *n* = 53) was tracked (2W: *M* = 3.93, *SD* = 1.39, ranging from 2 to 7; 6M: *M* = 3.88, *SD* = 1.31, ranging from 1 to 7; 12M: *M* = 4.02, *SD* = 1.29, ranging from 1 to 6). The average number of the trials used was as follows: 2W: *M* = 86.07, *SD* = 1.39, ranging from 83 to 88; 6M: *M* = 86.12, *SD* = 1.31, ranging from 83 to 89; 12M: *M* = 85.98, *SD* = 1.29, ranging from 84 to 89. To maintain comparability, an equal number of trials were randomly selected from each condition. ERP waveforms were baseline-corrected using the 200 ms pre-stimulus-onset window by substracting the mean amplitude of the baseline period from the amplitude at each time point of the ERP waveform.

As a quality control measure, grand average ERPs were calculated per subject at each developmental stage, comparing brain activity during the beep tone sequence to a no-stimulation condition of silence with no auditory stimulation. For each subject, the ERP waveform was visually inspected after averaging across trails to assess data quality. Participants with excessively noisy or non-interpretable waveforms were excluded from further analysis. Consequently, at each measurement point, exclusions based on data quality concerns were made (2W: *n* = 21; 6M: *n* = 8; 12M: *n* = 9). This resulted in the final sample size of *n* = 41 for 2W, *n* = 58 for 6M and *n* = 53 for 12M.

All ERPs plots shown in the results section were generated using R (Version 4.3.3; R Core Team, 2021) in RStudio (Version 2025.05.0+496; Posit Team, 2025).

#### 4.6.2. Inter-Trial Phase Coherence (ITPC) Analysis

ITPC analysis (2W: *n* = 41; 6M: *n* = 58; 12M: *n* = 53) was conducted using the same fronto-central electrodes, preprocessing pipeline, and segmentation of control (no-stimulation)-periods (2W: 8 to 28 s; 6M/12M: 10 to 160 s) as the event-related analysis described in Section *4.6.1*. Epoch rejection criteria also remained unchanged (2W: *M* = 4.51, *SD* = 2.12, ranging from 0 to 8; 6M: *M* = 4.28, *SD* = 2.25, ranging from 0 to 8; 12M: *M* = 4.34, *SD* = 2.32, ranging from 0 to 11). On average, the following number of trials was used for analysis: 2W: *M* = 85.49, *SD* = 2.12, ranging from 82 to 90; 6M: *M* = 85.72, *SD* = 2.25, ranging from 82 to 90; 12M: *M* = 85.66, *SD* = 2.32, ranging from 79 to 90. Epochs spanned -4 s to +4 s relative to stimulus onset. Complex Fourier spectra were computed using a multitaper convolution method. Phase angles were extracted from the spectra, and ITPC was calculated as the normalized vector sum of phase angles across trials. Baseline correction was applied by computing relative changes in the ITPC values with respect to the average ITPC in the 200 ms prestimulus-onset window.

#### 4.6.3. Time-Frequency (TF) Analysis

For time-frequency analysis the same data, electrode selection, and preprocessing steps as for the ERP and ITPC analysis were used. Epoch rejection, segmentation and trial matching procedures were identical to those used in the ITPC analysis. Time-frequency representations were computed using the same multitaper convolution method. Frequencies ranging from 0.5 to 30 Hz were analyzed in 0.5 Hz steps with a sliding window of 500 ms length, tapered by Hanning window. Power was computed every 500 ms over a time range of -4 s to 4 s relative to stimulus onset. Resulting power spectra were baseline-corrected using a relative change method, computed as the percentage change from the average power in the 200 ms prestimulus-onset window.

#### 4.6.4. Statistical Analyses

EEG data at three developmental stages (2W, 6M, 12M) were analyzed to investigate neural responses (ERP, ITPC, TF) both within each age group and across developmental stages. Within each stage, stimulation responses were compared to the no-stimulation condition. Longitudinal changes were assessed pairwise across developmental stages (2W vs. 6M, 2W vs. 12M, 6M vs. 12M) within subjects. Statistical analysis was performed using a cluster-based permutation test, as implemented in FieldTrip (Maris & Oostenveld, 2007). Paired-sample *t*-tests were conducted with a Monte Carlo correction using 5000 permutations (cluster-alpha = 0.05, critical-alpha = 0.025). We report cluster-level *p*-values and the sum of *t*-values (∑t) in the results section.

## Data and Code Availability

Data is available from the corresponding author upon reasonable request. The code used for the analyses in this study is available at https://github.com/evareisenberger/InfantEEGAnalysis.git

## Acknowledgement

We thank all mothers, fathers, and infants who participated in the study. Furthermore, we thank all students who helped and assisted with data collection. The authors also acknowledge the computational resources and services provided by Salzburg Collaborative Computing (SCC), funded by the Federal Ministry of Education, Science and Research (BMBWF) and the State of Salzburg.

## Declaration of Competing Interest

None.

## Funding

This work was funded by the Austrian Science Fund (FWF: 10.55776/P33630). C.F. and J.P. received financial support by the Doctoral College “Imaging the Mind” (FWF: 10.55776/W1233).

## CRediT Authorship Contribution Statement

**Eva Reisenberger**: Writing – original draft, Writing – review and editing, Formal analysis, Data curation, Project administration, Methodology. **Manuel Schabus**: Writing – review & editing, Resources, Methodology, Supervision, Funding acquisition. **Cristina Florea**: Writing – review & editing, Project Administration, Data Curation, Methodology. **Monika Angerer**: Writing – review & editing, Project administration, Funding acquisition, Supervision, Methodology. **Michaela Reimann-Ayiköz**: Writing – review & editing, Project Administration, Data Curation, Methodology. **Jasmin Preiß**: Writing – review & editing, Project Administration, Data Curation, Methodology. **Dietmar Roehm**: Writing – review & editing, Supervision, Methodology. **Dominik P. J. Heib**: Writing – review & editing, Methodology. **Claudius Fazelnia:** Methodology. **Mohamed S. Ameen**: Writing – review & editing, Writing – original draft, Supervision, Formal Analysis, Methodology, Conceptualization.

## Declaration of Generative AI Technologies in the Manuscript Preparation Process

During the preparation of this work the authors used Academic AI (University of Salzburg, GPT-5) and ChatGPT-5 in order to rephrase sentences and thereby improve readability and clarity, and to occasionally help with debugging issues the authors came across during preprocessing and analysis. After using these tools, the authors reviewed and edited the content as needed and take full responsibility for the content of the published article.

## Appendix A (Figures)

**Figure A1.**
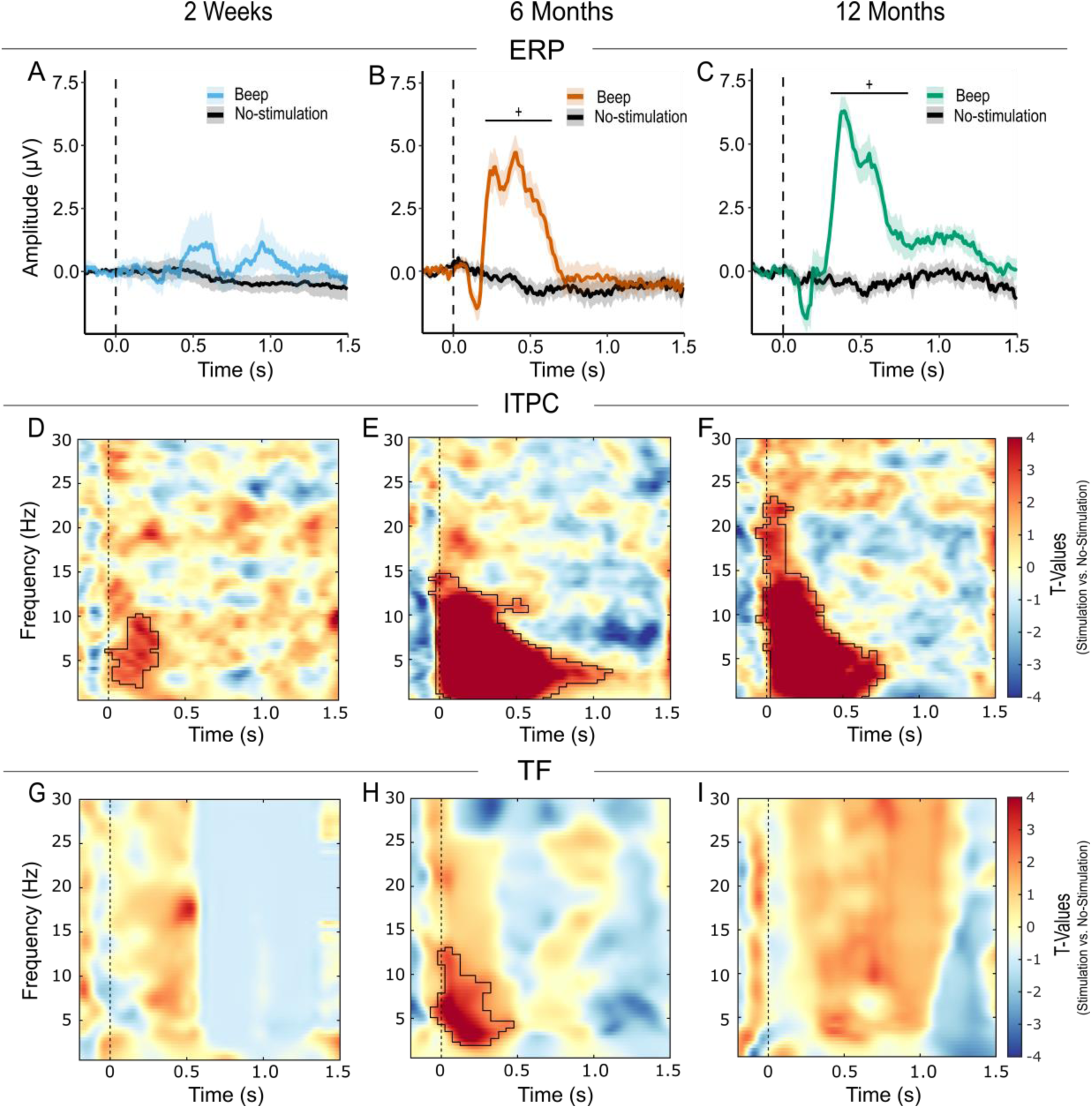
Overview of Auditory Neural Responses Within Age Groups Across Conditions (Stimulation vs. No-Stimulation) Recorded Over the Temporal Region. *Note.* Six electrodes from the temporal region were selected: E45 (T7) and E108 (T8) along with their immediate anterior and posterior neighbors (for E45: E39 and E50; for E108: E115 and E101). The signal reflects group level results from all participants available at each measurement time point (2W: *n* = 41, 17 female; 6M: *n* = 56, 28 female; 12M: *n* = 52, 26 female). Columns represent the three measurement time points. Each row displays a different type of analysis: **A-C**) Grand average event-related potential (ERP) waveforms for the stimulation condition at 2 Weeks (blue), 6 Months (orange), and 12 Months (teal). The no-stimulation control condition is plotted in black. Each panel displays mean ERP amplitude (µV) over time (s), aligned to stimulus onset (dashed vertical line at 0s). Shaded regions represent the standard error of the mean. Asterisks indicate statistical significance (\**p* < 0.05, ^+^*p* < 0.01) with the vertical line marking the corresponding time window. **D-F**) Time-frequency (TF) plots depicting t-values obtained through cluster-based permutation tests of inter-trial phase coherence (ITPC), representing the difference between stimulation and no-stimulation condition, and **G-I**) TF plots of spectral power differences shown as *t*-values from cluster-based permutation tests comparing the stimulation to the no-stimulation condition. In the second and third row, the TF plots display data across frequencies (0.5 to 30 Hz) and time (-0.2 s to 1.5 s), time-locked to the onset of the stimulus onset (vertical dashed line at 0s). The color scale represents *t*-values, where positive values (warm colors) indicate higher phase synchronization or spectral power during the presentation of the stimuli, and negative values (cool colors) indicate lower phase synchronization or spectral power relative to the no-stimulation condition. Significant clusters are outlined with a black contour within the TF plots.

**Figure A2.**
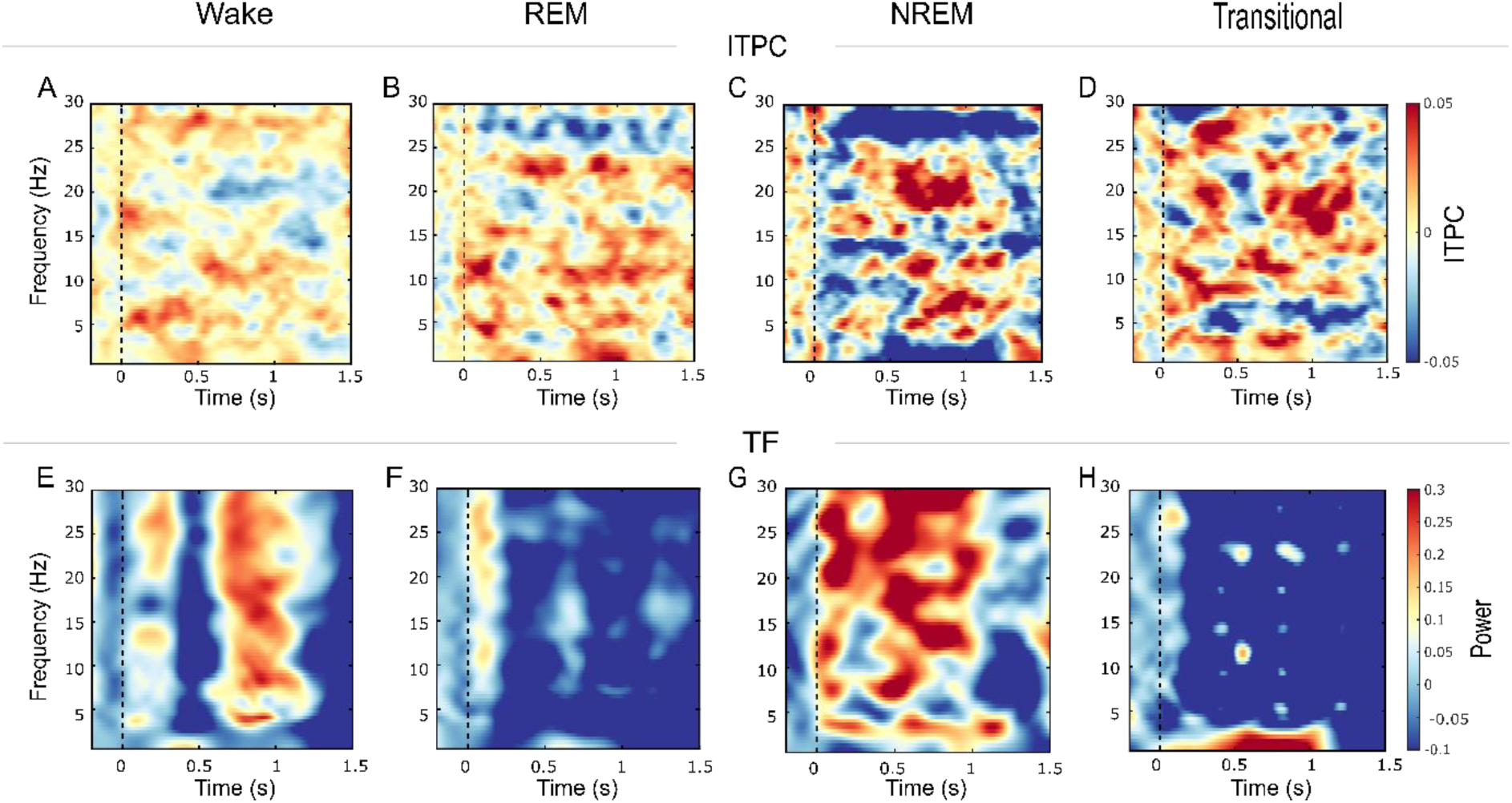
Additional Overview of Neural Responses at 2 Weeks After Birth Split by Sleep Stage. *Note*. Columns represent the four different sleep stages (*n* = 20 in Wake; *n* = 5* in REM; *n* = 2 in NREM; *n* = 6 in Transitional). Each row displays a different type of analysis: **A-D**) Inter-trial phase coherence (ITPC) difference plots (stimulation – no-stimulation) for each sleep stage. The color scale ranges from -0.05 to 0.05, representing changes in phase synchronization across trials. Warmer colors reflect increased phase synchronization during the stimulation compared to the no-stimulation condition, while cooler colors indicate decreased phase-locking during stimulation compared to no-stimulation respectively. **E-H**) Time-frequency (TF) power difference plots (stimulation – no-stimulation) separately for each sleep stage. The color scale ranges from -0.1 to 0.3, indicating changes in spectral power. Warmer colors represent increased power during stimulation compared to the no-stimulation condition, whereas cooler colors indicate spectral power decrease during stimulation compared to the control (no-stimulation) condition. *For oscillatory analyses in REM another participant was excluded due to substantial noise issues.

**Figure A3.**
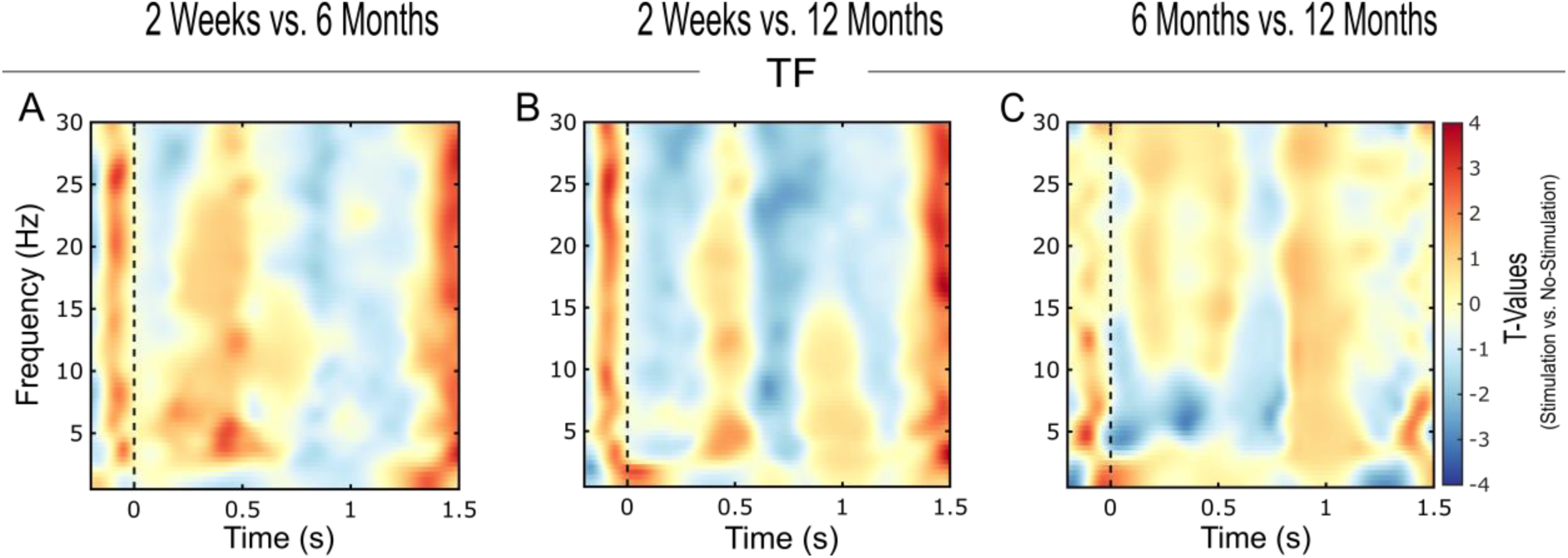
Pairwise Comparisons of Power Differences Between Developmental Time Points: A) 2 Weeks vs. 6 Months; B) 2 Weeks vs. 12 Months; C) 6 Months vs. 12 Months. *Note*. Columns represent the three pairwise contrasts: 2 Weeks vs. 6 Months (*n* = 17), 2 Weeks vs. 12 Months (*n* = 15) and 6 Months vs. 12 Months (*n* = 48). Each panel displays a time-frequency (TF) plot of power differences expressed as *t*-value statistics obtained via cluster-based permutation testing comparing the stimulation to the no-stimulation condition. The TF plots display data across frequencies (1 to 30 Hz) and time (-0.2 s to 1.5 s), time-locked to the onset of the stimulus onset (vertical dashed line at 0 s). The color scale for *t*-values represents statistical differences between timepoints, where positive values (warmer colors) indicate higher spectral power in the older age group, and negative values (cooler colors) indicating higher values in the younger age group respectively. No significant clusters appeared.

**Figure A4.**
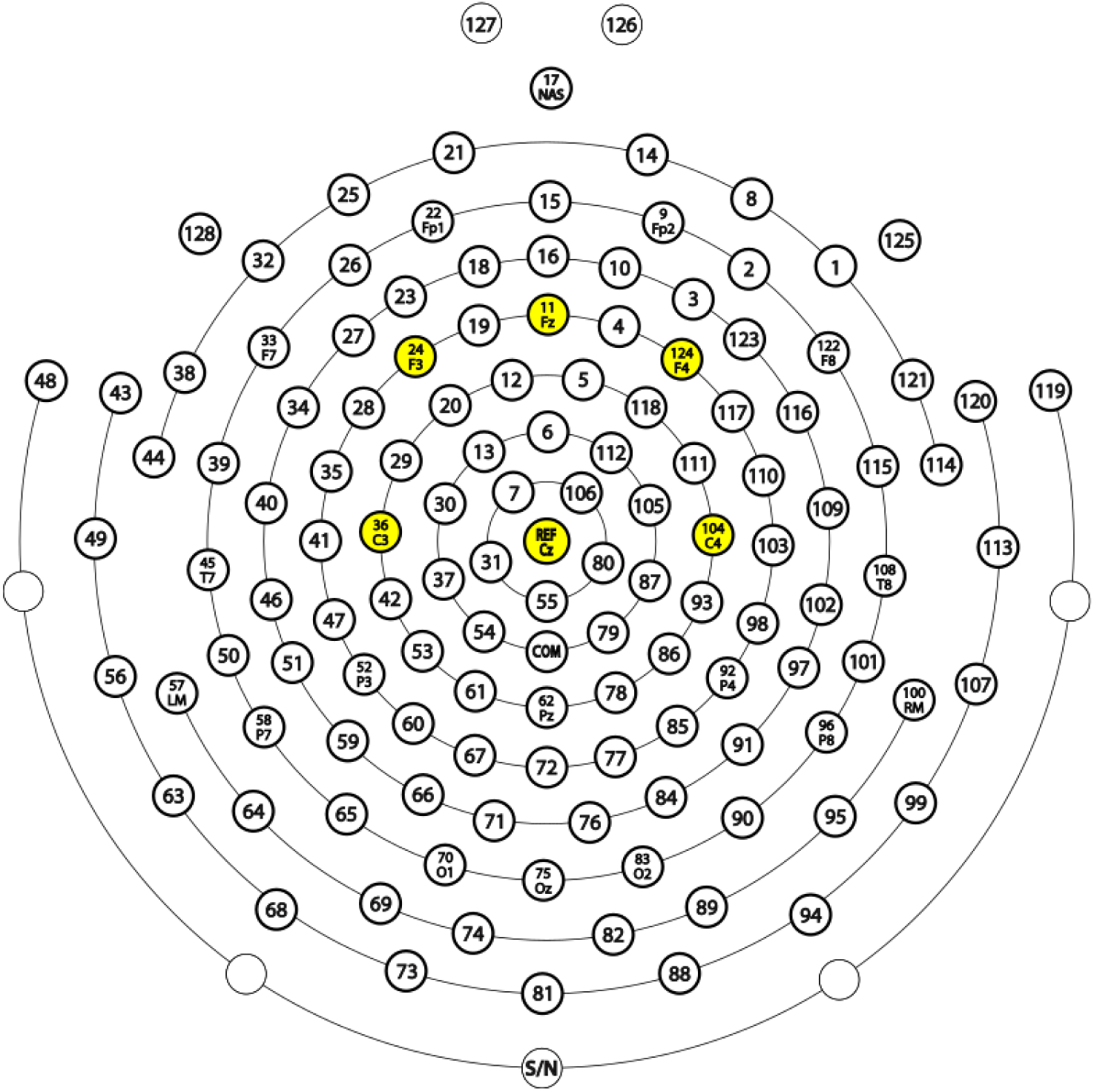
An Overview of the EEG Electrode Net Used. *Note.* In the main analyses, we focused on six electrodes located in the fronto-central region, which are highlighted in yellow in the figure. The EGI 128-channel EEG electrode layout is adapted from the Healthy Brain Network / FCON 1000 Project website with minor graphical modifications. The dataset for that resource is described in Alexander et al. (2017).

## Appendix B (Tables)

**Table B1.** Maternal Demographic and Family Characteristics of Participants Completing at Least One Postnatal Assessment (n = 69).

|  |  |
| --- | --- |
| Maternal Age at Study Entry, mean (sd) [range] | 32.9 y (4.72) [20-45] |
| *Family Status (%)* |  |
| Married | 47.8 |
| In a relationship | 46.4 |
| Single | 5.8 |
| *Highest School Qualification (%)* |  |
| High School Certificate | 85.5 |
| Vocational School | 14.5 |
| *Education (%)* |  |
| Apprenticeship | 18.8 |
| Vocational Secondary School | 20.3 |
| University Degree | 60.9 |
| *Parity (%)* |  |
| 1st | 71.0 |
| 2nd | 26.1 |
| 3rd | 1.45 |
| >5th | 1.45 |

**Table B2:** Birth and Neonatal Information of Participants Completing at Least One Postnatal Assessment (n = 69).

|  |  |
| --- | --- |
| Gestational Weeks at Delivery, mean (sd) [range] | 39+6 (8 d) [36+6-41+6] |
| *Mode of Delivery (%)* |  |
| Vaginal Delivery | 62.3 |
| Vaginal Operative Delivery | 10.1 |
| C-Section for Medical Reasons | 26.1 |
| C-Section for Personal Reasons | 1.45 |
| *Place of Birth (%)* |  |
| Delivery Room in a Hospital | 97.1 |
| Home | 1.45 |
| Other | 1.45 |
| Sex (% female) | 46.4 |
| Height, mean (sd) [range] | 51.7 cm (2.45) [48-58] |
| Weight, mean (sd) [range] | 3356 g (438) [2170-4740] |
| Apgar Score 10 min. after birth mean (sd) [range] | 9.89* (0.56) [6-10] |

*Note.* \*Apgar Scores were missing for three participants. Therefore, the presented statistics are based on *n* = 66 available values.

**Table B3.** Overview of Infants’ Positions During the Beep Tone Sequence at 2 Weeks, 6 Months and 12 Months, With Breastfed Infants Indicated Where Relevant.

|  |  |
| --- | --- |
| *2 Weeks (n = 41\*)* |  |
| Side-lying | 22 (14 breastfed) |
| Supine | 15 |
| Prone | 6 |
| *6 Months (n = 58)* |  |
| Side-lying | 2 (1 breastfed) |
| Supine | 8 |
| Prone | 2 |
| Sitting | 46 |
| *12 Months (n = 53)* |  |
| Side-lying | 1 (1 breastfed) |
| Sitting | 52 |

*Note.* \*Note that the total number of positions reported does not sum up to the reported number of , as infants s recordings ometimes changed positions during the beep tone sequence ( $n = 2$ out of 41).

## Notes

### Competing Interest Statement

The authors have declared no competing interest.

### Summary of Updates

Revised legend for Figure 4, typo corrections and minor layout adjustments.

